# Variation in pyrethroid resistance phenotypes in *Anopheles darlingi* from residual malaria transmission area: warning on suspected resistance built-up in French Guiana

**DOI:** 10.1101/2020.12.21.423491

**Authors:** Samuel Vezenegho, Romuald Carinci, Jean Issaly, Christophe Nguyen, Pascal Gaborit, Laetitia Ferraro, Guillaume Lacour, Emilie Mosnier, Vincent Pommier de Santi, Yanouk Epelboin, Romain Girod, Sébastien Briolant, Isabelle Dusfour

## Abstract

*Anopheles darlingi* is the major vector of malaria in South America. In French Guiana, malaria transmission occurs inland and along the rivers with a particular reemergence in the lower Oyapock area. Control against malaria vector is performed using deltamethrin indoor residual spraying and long lasting impregnated bednets. For four years, the level of resistance to pyrethroids was monitored using CDC bottle tests in *An. darlingi* populations. Resistance built-up was suspected in a mosquito population in malaria endemic area but did not sustained, supposably due to the reintroduction of susceptible alleles. No mutation on the insecticide target genes was found, metabolic resistance is then suspected.

## INTRODUCTION

*Anopheles darlingi* is the main vector of malaria in South America. For decades, this species has been targeted by vector control program using indoor residual spraying and bednets distributions. *Anopheles darlingi* has a high adaptability in its behavior depending on the locality in its distribution^1,2^. Known as a sylvatic mosquito, its density drastically increases in presence of human^3^. However, few evidence of insecticide resistance exists in that species^4,5^. This observation could be explained by the massive genetic exchange between the forest and the human dwellings where vector control is operated, that dilutes the resistance effect^6^.

In French Guiana, a French overseas territory, *An. darlingi* is found across the territory and remains strongly associated to malaria outbreaks. Those outbreaks and transmission of *Plasmodium vivax* and *P. falciparum* occur along the rivers and inland close to goldmines^7–10^. A drastic decrease of malaria incidence has been observed the last 20 years, but the area of Saint Georges de l'Oyapock, at the border with Amapa state (Brazil), remains impacted^11^. In areas with high malaria transmission, excluding remote area in the Guyanese forest, indoor residual spraying with deltamethrin is recurrently implemented. Since 2012, long lasting impregnated nets (LLINs) have been distributed to the population. Furthermore, spatial sprays of deltamethrin have recently been performed during malaria outbreaks. Additionally, French Armed Forces involved in illegal goldmining control use permethrin-impregnated uniforms^12^. Molecules from the pyrethroid family are the only one available and authorized for adulticide use under the European and French regulation. In a context of malaria control, means to reduce vector must be evaluated and insecticide efficacy monitored.

## MATERIAL, METHODS & RESULTS

Two populations of *An. darlingi* from French Guiana were studied. The first one was located at La Césarée (LC) farm in Macouria at 45-minutes’ drive west from Cayenne, the capital town. This equine and bovine farm is surrounded by a humid savannah. No insecticidal treatment is performed in that area. The second site was situated in Blondin (BL) hamlet, accessible by 5-minutes boat from Saint Georges de l'Oyapock and located on the Brazilian border to the east. The population living in this location along the Oyapock River suffered from recurrent malaria cases. IRS is implemented there each year from September to December to prevent malaria transmission. Inhabitants are equipped with LLINs impregnated with deltamethrin (2012) then alpha-cypermethrin (2016). Mosquito collections were performed in 2014, 2015, 2016 and 2018 using either supervised human landing collection (HLC) or subsequently mosquito magnet trap (MM) (Table 1) (especially in 2018, because of a malaria outbreak). Local volunteering residents were trained on HLC and informed of the associated risks. Malaria prophylaxis was proposed and information on the medication was provided. Collectors who benefited from prophylaxis gave their free, expressed and informed consent. CDC-bottle tests were performed either in our field lab atSaint Georges de l'Oyapock with freshly collected mosquitoes from Blondin or in our Cayenne main lab with mosquitoes captured at La Césarée. Sixty-seven to 316 mosquitoes were exposed to the insecticide per time point, site and insecticide along with control bottle for 4 to 12 replicates (Table 1). Insecticide resistance was evaluated for deltamethrin (12.5μg/bottle), permethrin (21.5μg/bottle) and alpha-cypermethrin (12.5μg/bottle). Additionally, malathion (50μg/bottle) was tested at Blondin in 2014 and 2015 (Table 1). The number of knocked-down mosquitoes in the impregnated bottle, which included both dead and unable to fly mosquitoes, were recorded every 5 to 15 min minutes until 100% of them were down (Table 1). Species identification was performed after the test. Raw data were corrected by Abbott's formula when needed.

**Table 1:**
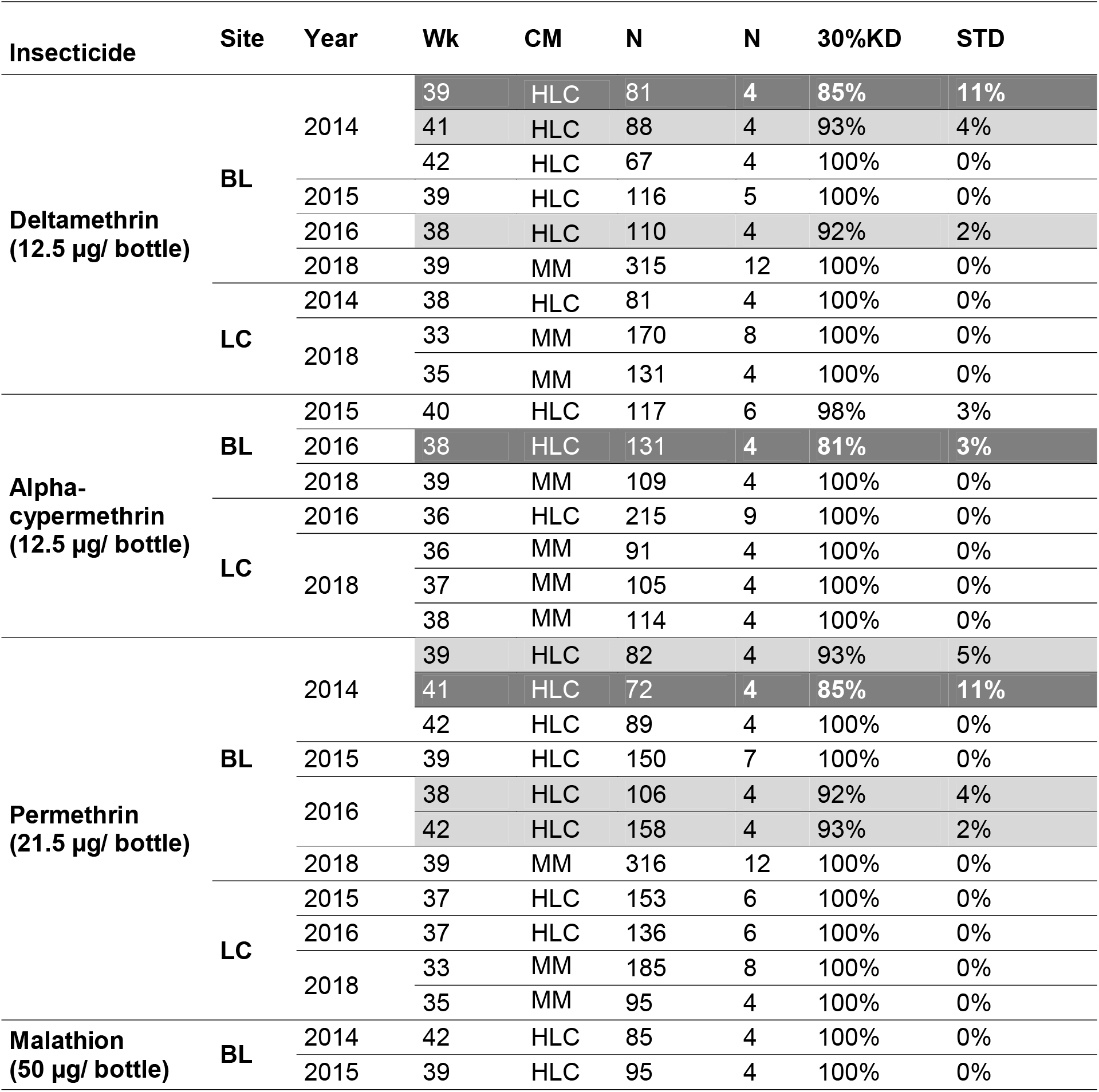
Percentages of knocked-down mosquitoes at 30 minutes (30%KD) and the standard deviation (STD) amongst replicates in CDC-bottle test for *Anopheles darlingi* populations collected in Blondin (BL) and La Césarée (LC). Test results are break down into the insecticidal molecule, year and week (Wk). In addition, the method of collections (CM) *i.e.* human landing catch (HLC) and mosquito magnet trap (MM), the number of mosquitoes tested (N) and replicates are mentioned (n). Dark grey: resistant populations with 30%KD < 90%; light grey: 98%<30%KD< 99% possible resistance; white: susceptible 30%KD>98%.

Therefore, we obtained knock-down rates at 30 min (30%KD), the recommended diagnostic time, and also a KT50/95 value which is the theoretical time to knock down/kill 50%/95% of mosquitoes. This time is calculated based on datasets with a minimum of four time-points producing from 0 to 100% of 30%KD. The probit regression of the knocked-down response over time was performed in BioRssays^13^ in R environment, which produces *p-value* for the logistic regression significance. The *p-values* equals to 0.97 and above permitted to consider the regression valid. The script also permits to compare regression results using chi-square test. In addition, the ratio of resistance RR50 or RR95 generally calculated with the KT values divided by the same values for reference strain were calculated per insecticide by dividing the obtained values by the minimal KT in the panel.

All mosquitoes were identified as *An. darlingi*. All populations with a 30%KD above 98% were diagnosed as susceptible according to the rules for CDC-bottle test interpretation. As expected, La Césarée population was susceptible to all pyrethroids whatever the date. Blondin population was susceptible to malathion with 100% of dead mosquitoes at 30 minutes in 2014 and 2015 (Table 1). However, we observed that 30%KD values varied below 90%, the threshold for resistance, up to 100% for pyrethroid compounds in Blondin (Table 1). In 2014 and 2016, those results had changed from one week to another from resistant to. Susceptibility was recorded in 2015 and 2018 for which only one week of test was performed.

To investigate further those fluctuations, KT50 and associated Resistance Ratio (RR) were obtained. First, they confirmed the trends observed for the end-point measures at 30 minutes. La Césarée population had the lowest KTs overall while KT from BL-2016-38 population where the highest in Blondin for all insecticides. The lowest values at la Césarée were used to calculate RR50 for each insecticide (Table 2, underlined values). The ratio calculated with those values highlighted that the population of 2014-BL-38 exposed to deltamethrin and that of BL-2016-38 exposed to any pyrethroids had resistance ratio (RR50) above 2, the threshold to diagnose resistance. Secondly, we observed that the slope of regression was lower in BL than in LC population, suggesting that even if the data do not support resistance, it takes more time to knock down mosquitoes with pyrethroids in BL than in LC population.

**Table 2:**
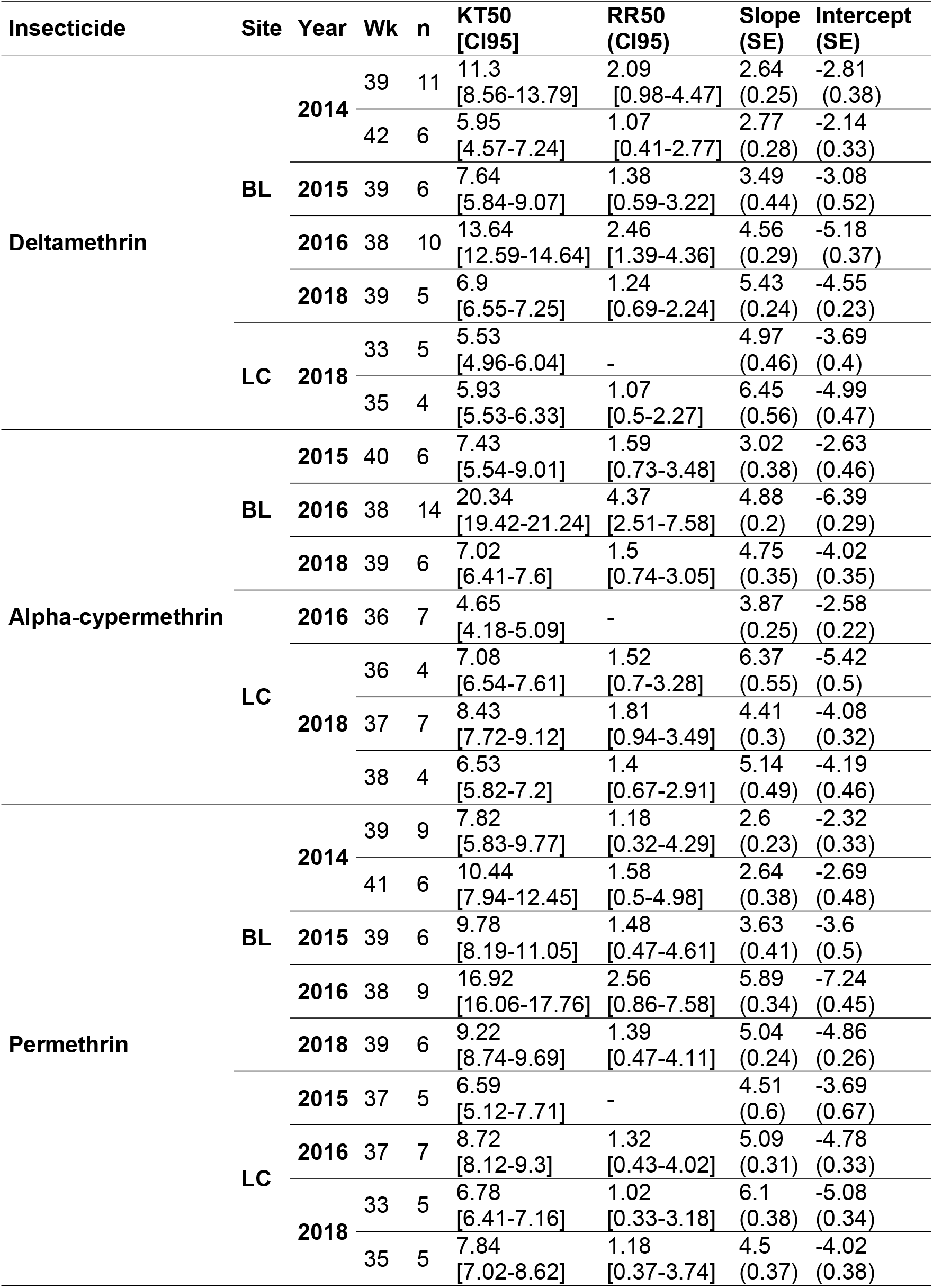
Results of significant regressions performed per pyrethroid molecule, site, year and week (Wk): knocked-down Time 50 (KT50), ratio of resistance (RR50) with their confidence interval at 95% (CI95), parameters of regression (slope and intercept with their standard errors) are represented. The number of time points per regression is also mentioned (n).

To complement the study, surviving (n = 16) and dead (n = 32) *An. darlingi* mosquitoes exposed to diagnostic concentrations of pyrethroid insecticides and malathion (caught at Blondin hamlet in 2014) were used to amplify and sequence a fragment of the acetylcholinesterase 1 (ACE1) gene as described in Weill (2004)^14^ and a fragment of the voltage-gated sodium channels (VGSC) gene as described in Orjuela (2019)^15^. This last one spanning codons 1010, 1013 and 1014 at the S6 segment of domain II to identify point mutations, which have been associated with insecticide resistance in different species of *Anopheles* malaria vectors. No mutation on the VGSC fragment nor ACE1 gene were detected in the 48 tested *An. darlingi*. Those sequences obtained 100% homology with amino-acids sequences already published (GenBank Accession numbers MN0503065 and AJ566402, for VGSC fragment and ACE1 gene respectively).

## DISCUSSION

The observations made over the years, rise the concern of insecticide resistance development in *An. darlingi* population in places were the use of pyrethroids is important^16^. In Blondin, both LLIN and IRS are performed with pyrethroids and *Aedes* control is performed occasionally as well. The population also uses household insecticides for several purposes.

In the recent years, IRS is not routinely performed anymore but only in case of outbreak along with spatial spraying, while distribution of LLINs is sustained by Public health authorities. In 2014, Blondin was the epicenter of malaria transmission in Saint Georges de l'Oyapock. After several years without large outbreaks, a drastic resurgence appeared in Trois Palétuviers in 2017. This hamlet is situated downstream of STG on the Oyapock river. In the meantime, French Guiana population suffered from chikungunya and Zika outbreaks against which pyrethroid-based control had been performed. None of those insecticidal applications are implemented in La Césarée.

Nevertheless, *An. darlingi* resistant populations are rare across its distribution on the opposite of *Aedes aegypti* for which a drastic increase of resistance was observed^17^. In fact, constant efficacy of control methods was already observed in the past in comparison with *Ae. aegypti* population that became resistant to DDT. The hypothesis formulated at that time was the continuous introduction of susceptible mosquitoes from forested areas to urbanized ones. Population genetic studies support this hypothesis^6^. *Anopheles darlingi* resistance development seems to be slowed down or nullified by a dilution effect. This factor would explain the variation in our monitoring.

Resistance to pyrethroids is a crucial point in French Guiana. As European territory, restrictive use of adulticide occurs. In consequence, only pyrethroids are available. This monotherapy conducts to select further resistance knowing that those molecules are also used in household sprays and against pest mosquitoes. In addition, LLIN providers have not renewed market released authorization for European markets, restraining the number of available products.

Insecticide resistance surveillance should be sustained in those places and a management plan set up. To reach this goal molecular markers of resistance should be obtained to early detect resistance. As shown in our study and a previous one in Colombia^15^, mutations on the sodium voltage gated gene is not involved in our *An. darlingi* populations. Further works on metabolic resistance are required as *An. darlingi* populations from Colombia exhibit metabolic resistance involving oxidases and esterases^18^. In addition, vector control method should be evaluated to maintain an efficient toolbox in a context of malaria control in the region.

## ACKNOWLEDGEMENTS

The authors want to thanks the people from Blondin and La Césarée, and the Camp Bernet staff of Saint-George’s town for their welcome and support in this study.

